# Temperature, but not excess of glycogen, regulate *post-mortem* AMPK activity in muscle of steer carcasses

**DOI:** 10.1101/2020.02.10.941666

**Authors:** P Strobel, A Galaz, F Villaroel-Espíndola, A Apaoblaza, JC Slebe, N Jerez-Timaure, C Gallo, A Ramírez-Reveco

## Abstract

*Post-mortem* muscle temperature affects the rate of decline in pH in a linear manner from 37.5 °C down near 0 °C, and this pH decline is correlated with the enzymatic degradation of glycogen to lactate. This transformation occurs in an anaerobic context that includes the metabolic splice between glycogenolysis and glycolysis; and both processes are strongly upregulated by AMPK enzyme. In this study we reported changes (0.5 h and 24 h post-mortem) in muscle glycogen concentration, lactate and AMPK activity from 12 samples of *Longissimus dorsi* from 38 steers that produced high pH (>5.9) and normal pH (<5.8) carcasses at 24 h post-mortem. Moreover, we evaluated changes in AMPK activity in samples from both categories incubated at 37, 25, 17 and 5 °C and supplemented with exogenous glycogen. Finally, we analysed if there were structural differences between polymers from both categories. Our analyses show that enzymatic AMPK activity was significantly higher at 17 °C than at 37 °C or 25 °C (p<0.0001 and p<0.05 in samples from normal and high pH categories, respectively), and was near zero at 5 °C. On the other hand, AMPK activity did not change in relation with excess glycogen and we did not detect structural differences in the polymers present in samples from both categories. We concluded that post-mortem AMPK activity level is highly sensitive to temperature and not at in vitro changes in glycogen concentration. Their results suggest that that normal levels of *pre-mortem* muscle glycogen and an adequate cooling managing of carcasses are relevant to let an efficient glycogenolytic/glycolytic flow required for lactate accumulation and pH decline, trough of *post-mortem* AMPK signalling pathway.

## Introduction

The enzyme that “reads” the energy level in the cellular system is the adenosine monophosphate (AMP)-activated protein kinase (AMPK). This enzyme is a heterotrimeric complex formed by the α (catalytic), β (regulatory) and γ (binding to nucleotides) subunits (Hardie et al., 2006). Homologous genes of these three subunits are present in all eukaryotic species with sequenced genomes (Hardie et al, 2003). The main function described for AMPK is the coordination of metabolic processes (anabolic and catabolic) in various tissues including cardiac and skeletal muscle, adipose tissue, pancreas, liver and brain tissues (Kahn et al, 2005; Ramamurthy and Ronnett, 2006). AMPK enzyme responds to hormonal, physiological and pathological stimulus, thereby inhibiting adenosine triphosphate (ATP) consumption (anabolic process), and but also activates ATP production (catabolic processes). Hence, consequences of AMPK activation is accompanied by an acute regulation of energy metabolism and chronic changes in gene expression. These changes are accompanied by phosphorylation of several effector proteins that include biosynthetic enzymes, transporters, transcription factors, ion channels and signaling proteins of the cell cycle (Leff, 2003; Hardie, 2004; Hallows, 2005).

The two isoforms of AMPK catalytic subunits (α1 and α2) are activated by an increase in AMP levels and a fall in ATP levels (e.g. exercise and stress) (Lizcano et al., 2004). AMPK is also activated by metformin/phenformin, drugs commonly used in the treatment of type II diabetes (Zhou et al., 2001). In all eukaryotes, the glycogen-binding domain (GBD) contained in beta subunits, causes AMPK complexes to associate with glycogen in cultured cells and cell-free systems (Hudson et al., 2003; Polekhina et al., 2003).

At a physiological level, nerve and muscle functions in most vertebrates are seriously affected by freezing, to which three stress components have been associated: changes in the cellular ionic, ischemia, and low temperature (Storey and Storey, 2004) (Dalo et al, 1995; Hillman, 1988, Lutz and Reiners, 1997). Environmental stress can also modify AMPK activity. Hypoxia and hypothermia in frogs increase the AMPK phosphorylation level and AMPK activity, measured by elongation factor 2 (EF2) phosphorylation (Bartrons et al., 2004). Effectively, winter survival of hundreds of species depends on freeze tolerance and one of these species (*Rana sylvatica*) revealed that during in vivo freezing, AMPK enzyme increases its activity (Storey, 2006). This suggests that AMPK activation (as a key modulator) would facilitate the best adaptation to hypothermia, freezing and thawing.

There are many studies that correlate glycogen content with the regulation of AMPK, although the underlying molecular mechanisms are poorly understood and some paradoxes remain (McBride et al, 2009). For example, while Polekhina et al. (2003) did not detect an effect of glycogen on the activity of purified rat liver AMPK; Wojtaszewski et al., 2003 found that the glycogen loading of skeletal muscle suppresses AMPK activation by exercise in humans or by contraction or 5-Aminoimidazole-4-carboxamide ribonucleotide (AICAR) administration in rodents (Derave et al., 2000; Wojtaszewski et al., 2002). Others have stated that AMPK is hyperactivated by exercise in subjects that accumulated excess glycogen, derived from McArdle disease, an inherited defect in glycogen phosphorylase (Nielsen et al., 2002). In the opposite direction, AMPK has a role in the regulation of glycogen synthesis. Muscle glycogen synthase (mGS) is phosphorylated at site 2 (Ser-7) by AMPK, which inactivates mGS (Carling and Hardie, 1989; Hardie and Sakamoto, 2006); however, AMPK promotes glucose uptake and glycogen accumulation in muscle (Wojtaszewski et al., 2002). Phosphorylation of mGS by AMPK causes a decrease in activity at low concentrations of the allosteric activator glucose-6-phosphate (Skurat et al., 1994). These phosphorylation events induced by AICAR treatment in muscle, occur only in wild-type mice, but not in AMPK-α2 knockout mice, showing that this site is a physiological target for AMPK (Jorgensen et al., 2004).

Regarding AMPK/Glycogen interaction, a study from McBride et al., (2009) showed that glycogen inhibits purified AMPK in cell-free assays, by direct action of GBD and that it varies according to the branching content of the glycogen. Moreover, these authors showed that the oligosaccharides used, are allosteric inhibitors of AMPK that also inhibit phosphorylation and activation by upstream kinases. This suggests that the GBD is a regulatory domain that allows AMPK to act as a glycogen sensor in vivo. More recently, it has been demonstrated that glycogen regulation occurs both in the presence and absence of AMP and is dependent on its binding to the carbohydrate-binding loop of the carbohydrate-binding module (CBM) (Scott et al, 2014).

Some studies have clearly shown that in *post-mortem* muscles there is a strong correlation between AMPK activation and pH decline, indicating that AMPK regulates glycolysis in *post-mortem* muscles (Du et al 2007; Shen & Du, 2005; Shen et al., 2006). In a murine genetic model, muscle samples from AMPK knockout mice, whether having exercised or not, showed higher ultimate pH, than samples from wild type mice (Shen and Du, 2005). In porcine *longissimus* muscle, an earlier and faster activation of AMPK is responsible for lower pH and higher lactic acid levels in PSE (pale, soft exudative) meat (Shen et al, 2006). In bovine *longissimus* muscle, we showed that AMPK activity was four time higher in carcasses that experienced a normal decrease in pH at 24 hours. This decline was conditioned by the reserve levels of initial glycogen at moment of slaughter and after the glycogenolytic/glycolytic flow in the L. dorsi muscle (Apaoblaza et al, 2015). In the same context, there are insufficient data about the changes in *post-mortem* muscle AMPK activity associated to low temperature decline. The aim of this study was to determine changes in *post-mortem* AMPK activity due to temperature decline and excess of glycogen level in muscle samples from steer carcasses with normal final pH (<5.8) and high final pH (>5.9).

## Material and methods

### Animals, slaughter, collection, selection, processing and pH categorization of samples

A total of 12 samples were sub-selected from a group of 38 carcasses previously categorised, selected and analysed in the study of Apaoblaza et al (2015). Selected steers were of similar age, had been under the same fattening system and had similar live weights and fatness. They had been transported for 4 h in a single group in the same truck to the slaughterhouse where they were kept in a group, had a pre-slaughter lair age time of 16 h and were all slaughtered on the same day and time. All carcasses were kept in the same cold chamber (T° between 0 and 4 °C). Samples of approx. 1 g were taken from the *Longissimus dorsi* (LD) muscle of the carcasses (at the level of the 10th rib) and were obtained using a Bergström biopsy needle at 0.5 h *post-mortem* (T0). These samples were placed in Eppendorf tubes, identified and immediately frozen in liquid nitrogen (−196 °C) and maintained in an ultra-freezer (−80 °C). After 24 h of chilling (0–4 °C), the final temperature (T24) and pH (pH 24) were measured, and a second muscle sample was taken from all carcasses at the same place. Six samples that had a pH 24 N 5.9 (high pH, n=6) and six samples that had a pH 24 b 5.8 (normal pH, n =6) were selected for this study.

### Temperature kinetic in carcasses

To establish the kinetic decline of temperature and pH (initial and final) of the carcasses in the cold chamber from T0 to T24 hours, a thermocouple/ electrode penetration probe with data logger (PCE 228 M, Spain) was connected to each carcass.

### Muscle glycogen concentration (MGC) determination

This analysis considered a glycogen purification and enzyme digestion method to obtain glucose (Chan and Exton, 1976). Briefly, 400 μl of 30% KOH was added to 100 mg of tissue, followed by incubation at 100 °C for 15 min with agitation. A 50 μl sample of each homogenate was placed on a 1 × 1 cm Whatman 31 ET filter paper. A calibration curve with glycogen (2.4 mg/ml) was prepared. Glycogen was precipitated with 66% cold ethanol for 10 min while stirring. Then, two washes were performed with 66% cold ethanol and the papers were dried to remove ethanol. Afterwards, 1 ml of amyloglucosidase (Sigma, 10113) solution 0.5 mg/ml in 400 mM sodium acetate buffer pH 4.8 was added and was incubated for 2 h at 37 °C in a stirred thermoregulated bath. Afterwards, 50 μl was extracted from each tube for the determination of glucose using the Glucose PAP liquiform kit (Labtest, cat. n. 84) and spectrophotometric assay based glucose oxidase and Trinder reaction, in which the product is a quinoneimine dye, which absorbs maximally at 505 nm (Yasmineh, Caspers, & Theologides, 1992). Finally, it was taken to initial glycogen concentration through stoichiometry.

### Lactate (LA) determination

Lactate concentration was determined by using the lactate oxidase method, liquiform lactate (Labtest Ref. 116). Briefly, muscle samples frozen in liquid nitrogen were homogenized in 20 mM citrate–50 mM phosphate buffer, EDTA 2.5mg/ml pH 6.8. An aliquot of the homogenate was used to determine the concentration of LA.

### Protein homogenization and AMPK activity assay

From each sample, 200 mg of muscle was powdered (liquid nitrogen) and homogenized in Ultra-Turrax T-10 (IKA-Werke GMBH, Germany) with an ice-cold buffer containing 500 μl lysis solution, 137 mM NaCl, 1 mM MgCl2,1% NP-40, 10% glycerol, 2 mM PMSF, 10 mM sodium pyrophosphate, 2.5 mM EDTA, 10 μg/ml Aprotinin, 10 μg g/ml Leupotinin and100 nM NaF. The homogenate was centrifuged at 20,000 g for 10 minutes; the supernatant was retained and stored at 80 °C. Protein concentration was determined by Bradford method (1976) using a commercial kit (Bio-Rad Laboratories; Hercules, CA, USA). The AMPK activity of homogenate muscle samples was determined by using the kinase assay Z-Lyte-Ser/thr 23 peptide (Life Technologies, PV4644), according to Apaoblaza et al (2015), with modifications. The employed assay is validated for AMPK (α1/2-β1-γ1) and based on fluorescence resonance energy transfer (FRET). A 20 μl peptide substrate Z-lyte 2X/ATP 2x in 5x kinase buffer was added to 20 μl sample diluted in 50 mM 10 mM MgCl Hepes buffer pH 7.5. The solution was incubated for 1 h at room temperature (initial and final characterization of the samples). Subsequently, 20 μl of Z-Lyte development solution was added, and samples were incubated for 1 h at room temperature. Finally, 20 μl of stop reagent was added to each well and then the signals of coumarin and fluorescein emission (445 and 520 nm respectively) were measured using the Thermo Scientific Varioskan Flash reader.

### Effect of temperature on AMPK activity

Similar to that described above, for this experiment, treatments consisted in incubation of the samples (both high and low final pH) for 1h at four different temperatures (5 °C, 17 °C, 25 °C and 37 °C) before addition of the stop agent and measure of fluorescein.

### Effect of glycogen addition on AMPK activity

Similar to that described above, for this experiment, treatments consisted incubation of the samples (both high and low final pH) for 1h at 17 °C with previous addition or not (control) with 60 mmol/kg, before to addition of the stop agent and measure of fluorescein.

### Glycogen analysis

For this analysis, homogenates were processed according to Villarroel-Espíndola et al. (2016). For the structural study of muscle glycogen from the samples, the spectrophotometric assay described by Krishman (1962) was used. Briefly, Iodine absorption spectra of glycogen are present in the homogenates of both category and time. Total glycogen was isolated from the respective samples by ethanol precipitation before spectrophotometric analysis. As a control, commercial glucose polymers was used, such as muscle glycogen and purified amylopectin.

### Statistical analysis

Statistical analyses were performed using one-way ANOVA with post hoc Bonferroni multiple comparison tests. We used GraphPAD (Software for Science) for analysis, and differences were considered significant and highly significant for P values of <0.05 and <0.01, respectively.

## Results

### Changes in pH, metabolic measure and AMPK activity between categories

Changes in muscular pH, glycogen content, lactate and AMPK activity at 0.5 and 24 hours in carcasses from both pH categories are shown in Table 1. Regarding final pH, the differences between high and low pH categories were closer to 0.71 pH point (6.44±0.05 vs 5.73±0.03, respectively). Consistent with the final pH, the differences in lactate accumulation at 24 h between high and low pH category were 47% (39.00±0.71 vs 57.43±2.15 mmol/Kg, respectively). Differences in muscle glycogen content present at 0.5 h, a main parameter used to define the pH evolution between high and low pH category were 365% (16.55±0.87 vs 76.94±6.9 mmol/Kg, respectively). Finally, like previous studies (Apaoblaza et al, 2015), the AMPK activity was higher in the samples categorized as normal final pH than high pH at both 0.5 and at 24 h (680 and 400%, respectively). All data are shown in Table 1, specifically AMPK activity, corresponding to the analyses carried out at room temperature (17-20 °C).

**Table 1:**
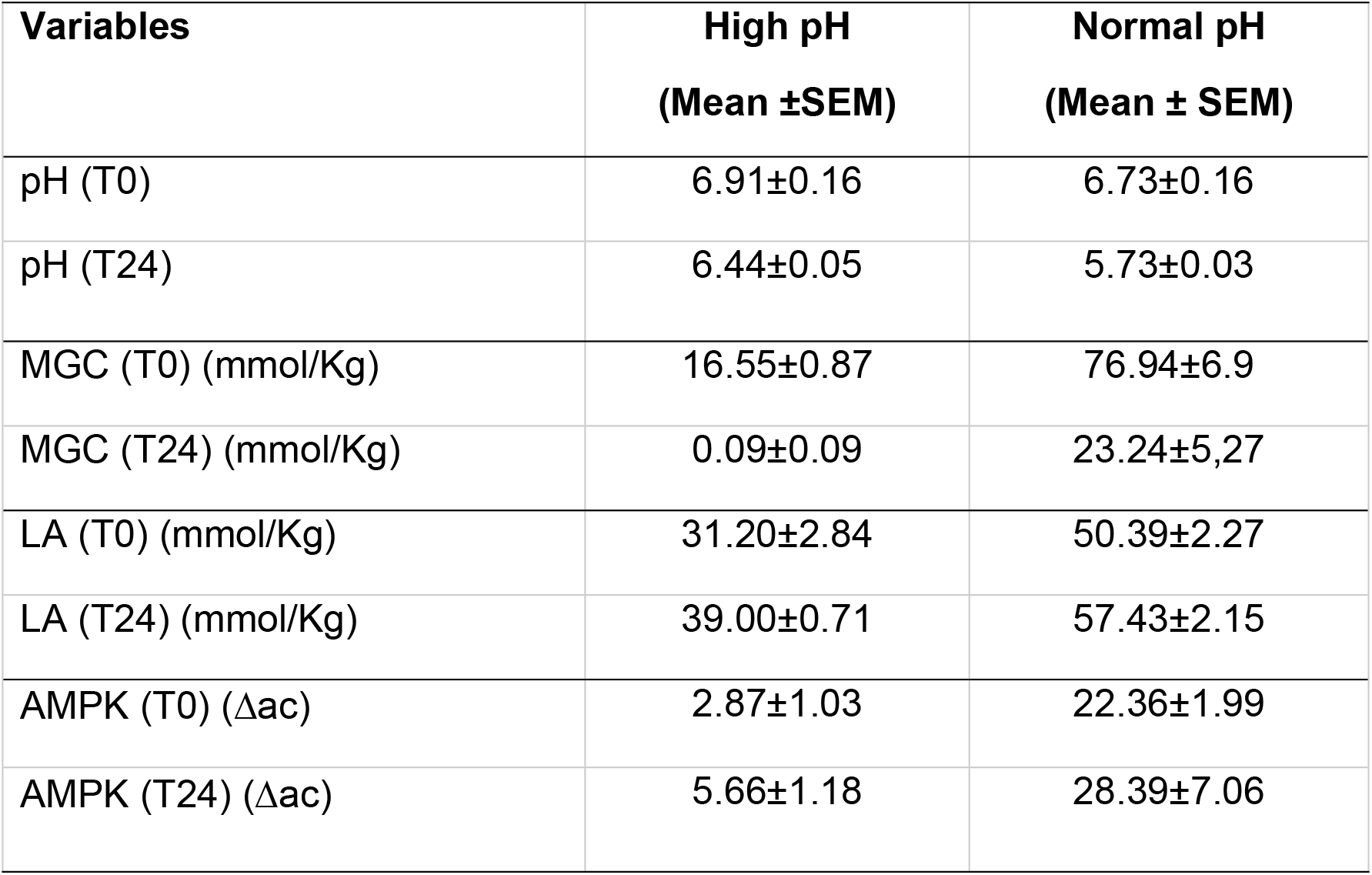
Mean pH, muscle glycogen concentration (MGC), lactate (LA) and adenosine monophosphate kinase (AMPK) activities in M. *longissimus dorsi*, measured at 0.5 h (T0) and 24 h (T24) *post-mortem* in steer carcasses categorized with normal (<5.8) v/s high (>5,9) final pH.

| Variables | High pH<br>(Mean ± SEM) | Normal pH<br>(Mean ± SEM) |
| --- | --- | --- |
| pH (T0) | 6.91±0.16 | 6.73±0.16 |
| pH (T24) | 6.44±0.05 | 5.73±0.03 |
| MGC (T0) (mmol/Kg) | 16.55±0.87 | 76.94±6.9 |
| MGC (T24) (mmol/Kg) | 0.09±0.09 | 23.24±5,27 |
| LA (T0) (mmol/Kg) | 31.20±2.84 | 50.39±2.27 |
| LA (T24) (mmol/Kg) | 39.00±0.71 | 57.43±2.15 |
| AMPK (T0) (Δac) | 2.87±1.03 | 22.36±1.99 |
| AMPK (T24) (Δac) | 5.66±1.18 | 28.39±7.06 |

### Temperature decline and changes in AMPK activity

The cooling rate in the cold chamber of the total carcasses used is shown in Figure 1. Analysis of the obtained non-linear function identified two components (slopes), the first from 32 to 10 °C (S1: −0.05 °C/min) and second from 10 to 2 °C (S2: −0.01 °C/min). Analyses of enzymatic activity of AMPK consistently showed a greater and sustained AMPK activity in muscle samples from carcasses categorized with normal final pH vs. those with high pH. Values were 680 and 400% at 0.5 and at 24 h, respectively (Table 1).

**Figure 1:**
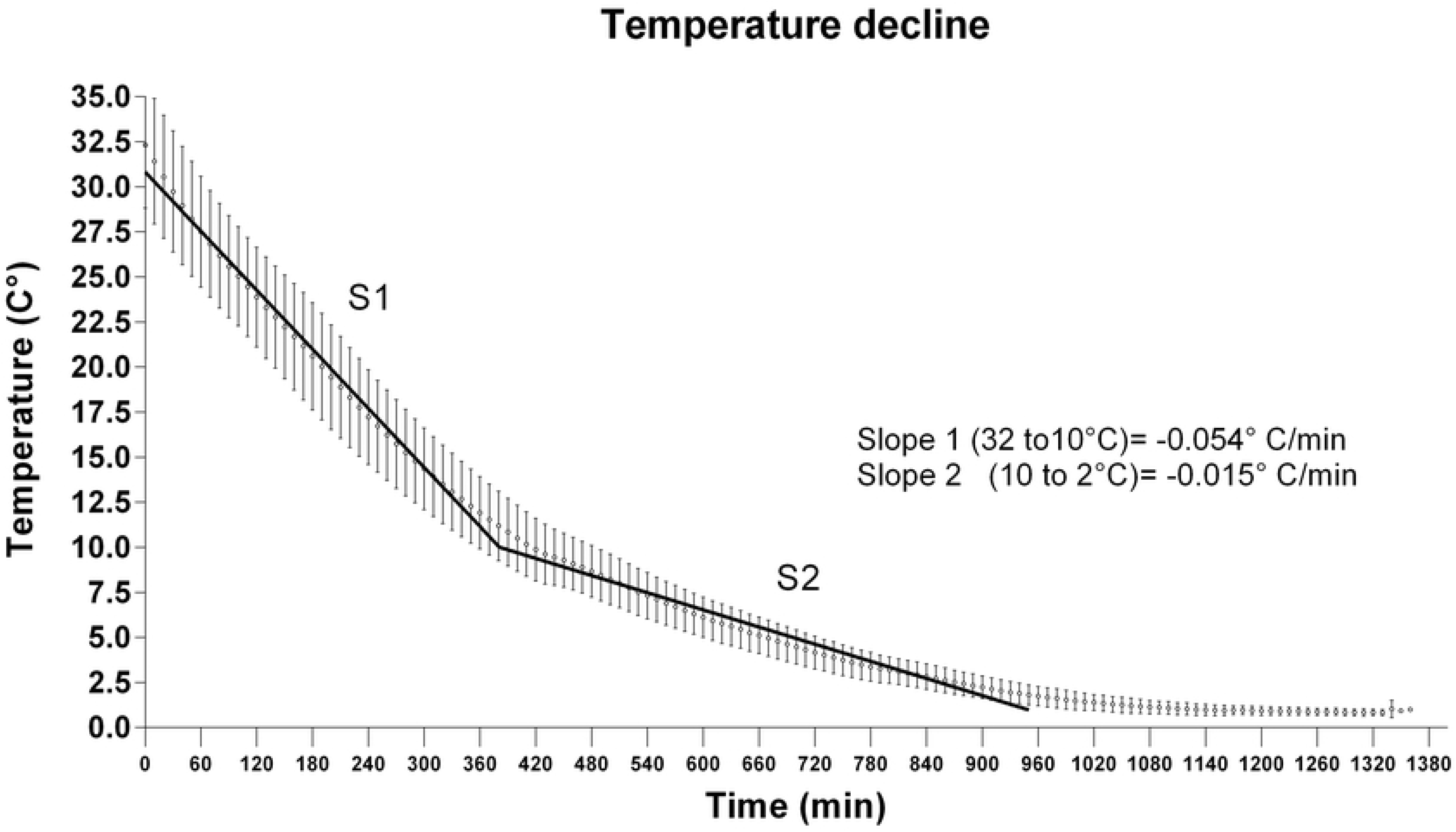
Temperature decline *post-mortem* in steer carcasses *M. longissimus dorsi* (Both categories, n=12). In the figure, slopes are shown (S1 and S2) corresponding to two components founded by non-linear function analysis.

Since enzymatic activity assays were performed at room temperature (approximately 17-20°C) and considering carcasses underwent significant temperature variations in the *post-mortem* “cooling rate” imposed on cold chamber, we further performed the AMPK activity analysis under three temperature points: 37 °C, 25 °C and 17 °C. Results from this experiment are shown in the Figure 2.

**Figure 2:**
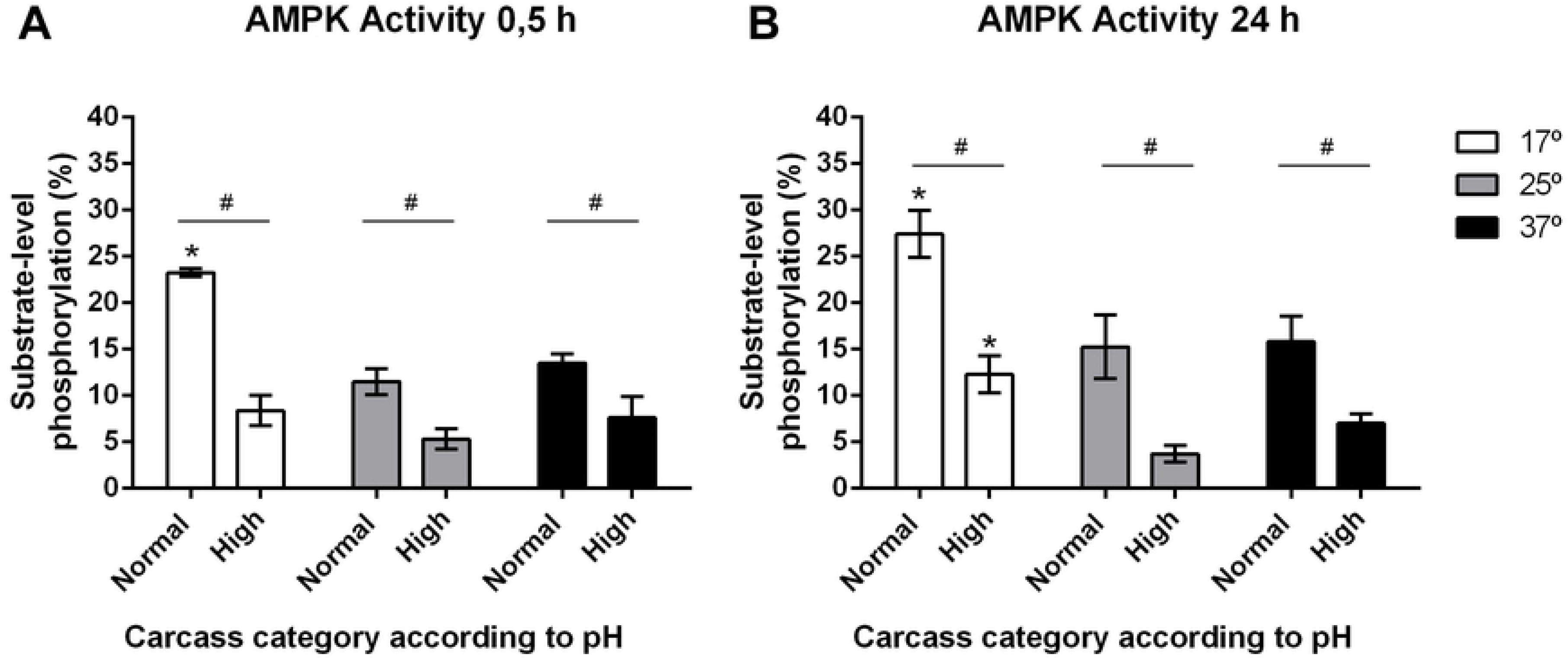
Average values ±SEM of AMPK enzyme activity levels in samples of in *M. longissimus dorsi* categorized with normal (<5.8) v/s high (>5,9) final pH, evaluated at 0.5 h and 24 h and in function of temperature. A: measure at 17, 25 and 37 °C at the 0.5 h; B: measure at 17, 25 and 37 °C at the 24 h. Two-way ANOVA and Bonferroni post-tests. Significant differences between categories (#, p<0.0001) and between 17 °C and 25 °C or 37 °C, within categories (*, p<0.0001), n=6.

Analysis of samples obtained from star cooling (0.5 h) and at three temperature analysed confirmed a higher AMPK activity (p <0.0001, n = 6) in samples taken at 0.5 h post-mortem, from carcasses that reached a normal final pH than those with high final pH (Figure 2A).

In general, AMPK activity at 0.5 h *post-mortem* did not vary according to temperature. The only exception was the analysis at 17 °C (samples with normal final pH), which showed to be significantly greater (p <0.0001) than at 25° and 37° (Figure 3A), and even higher (p <0.0001) with respect to measurements obtained at 17°, 25° and 37° in samples with high final pH (Figure 2A).

**Figure 3:**
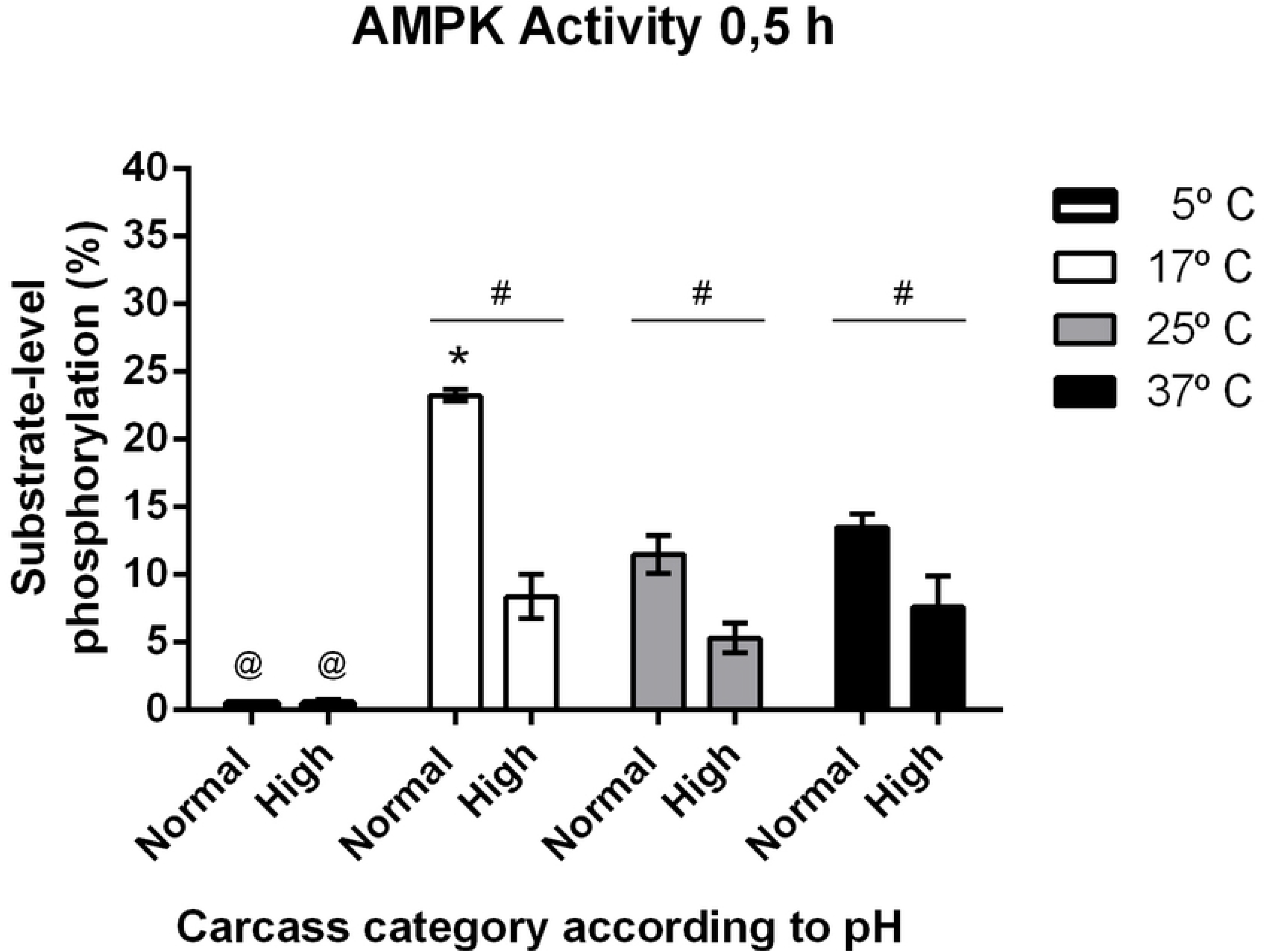
Average values ±SEM of AMPK enzyme activity levels in samples of in *M. longissimus dorsi* categorized with normal (<5.8) v/s high (>5,9) final pH evaluated at different temperatures in the cooling phase (5°, 17°, 25° and 37 °C at the 0.5 h). Two-way ANOVA and Bonferroni post-tests. Significant differences between categories (#, p<0.0001); between 17 °C and 25 °C or 37 °C, within categories (*, p<0.0001); between 17 °C and 5 °C, within categories (@. p<0.0001 and p<0.05, for normal and high final pH, respectively), n=6.

Comparative measurements at 24 h showed the same tendency, although less accentuated, with the novelty that average AMPK activity at 17 °C was significantly higher (p <0.0001, n = 6) in the samples from carcasses with high pH than those measurements made at 25 and 37°C, in the same category (p <0.05, n = 6). Figure 2B.

To expand the study of the effect of temperature on enzymatic activity of AMPK, a second experiment was performed with the inclusion of a thermal point closer to the final temperature of maintenance in the chamber (5°C). According to our records, this temperature is reached between 11-12 hours after the carcasses entered the cooling chamber (720 min on the x-axis, Figure 1). Considering that there were no significant differences associated with time (0.5 vs 24 h), only samples from biopsies taken at 0.5 h were analysed. The analysis confirmed that the activity of AMPK presented a strong regulation of temperature, showing maximal activity in the muscle samples with normal final pH, at 17° C (p <0.0001, n = 6); activity of AMPK was completely inhibited at 5° C in samples from both categories (p <0.0001, in normal final pH and p <0.05 in high final pH) (Figure 3).

### AMPK activity and glycogen content in post-mortem muscle

In order to evaluate the possible regulation of AMPK activity by available glycogen levels, an experiment was performed in which muscle samples from both categories were supplemented with 60 mmol / kg of glycogen (corresponding to the differences in initial MGC levels between samples from normal and high final pH). The AMPK activity for this study was measured at 17° C (temperature with higher AMPK activity) in samples from biopsies obtained at 0.5 h.

Under these conditions, glycogen levels reached by samples from both categories were the following: 76 (16 + 60) and 136 (76 + 60) mmol/kg of glycogen in the muscle samples with high and normal final pH, respectively. Results showed that the enzymatic activity of AMPK did not present changes associated with exogenous glycogen levels in any of the categories (Figure 4).

**Figure 4:**
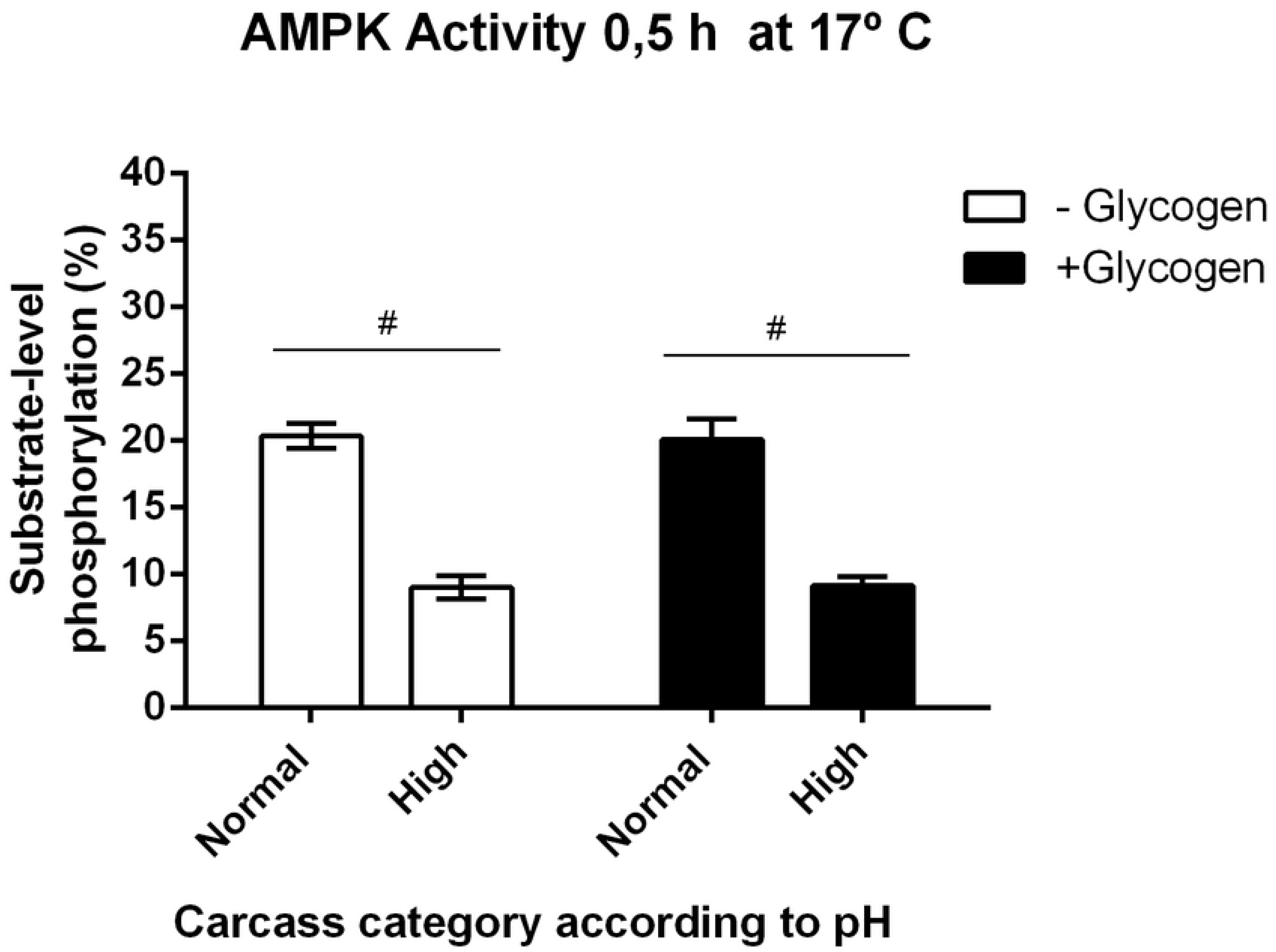
Average values ±SEM of AMPK enzyme activity levels in samples of *M. longissimus dorsi* categorized with normal (<5.8) v/s high (>5,9) final pH, evaluated at 0.5 h and to 17 °C, and in function of endogen and exogenous glycogen content. Two-way ANOVA and Bonferroni post-tests. Significant differences between categories (#, p<0.0001), n=6.

### Analysis of glycogen type

In order to evaluate if the glycogen type of samples present structural differences, an analysis of the absorption spectrum of glycogen in the presence of iodine was carried out (Figure 5A and 5B). Analyses showed that there was no difference between the means detected for glycogen extracted of samples from carcasses with normal and high final pH. Moreover, the obtained wavelengths coincided with those described for muscle glycogen molecules in the presence of potassium iodide, confirming that there were no changes in the three-dimensional configuration of the glycogen molecule, keeping a stable degree of branching in both conditions. This was evident when comparing the absorption spectrum of muscle glycogen purified by precipitation and the absorption profile characteristics of amylopectin, which have a lower degree of branching and have a displacement to the right of the maximum lambda, around 550 nm in Figure 5A.

**Figure 5.**
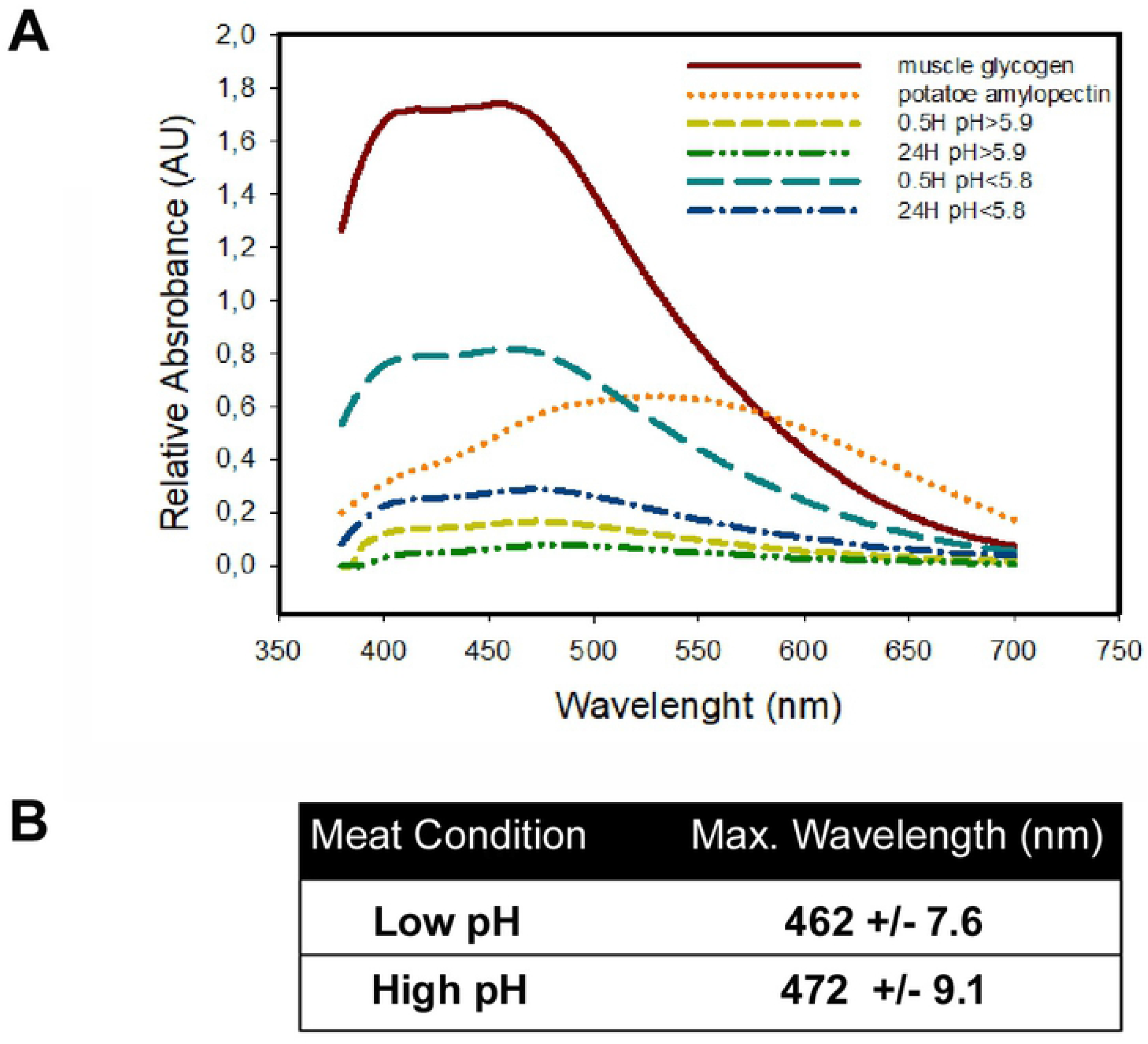
Structural glycogen analysis. A: Absorption spectrum of glycogen in samples of *M. longissimus dorsi* categorized with normal (<5.8) v/s high (>5,9) final pH, in presence of potassium iodide. B: maxima wavelength (nm). (n=6)

The spectrum analysis of glycogen, confirmed that the muscle samples collected from carcasses with high final pH, presented lower amounts of glycogen (in both 0.5 or 24 h), than those samples obtained from carcasses categorized with normal final pH (at both 0.5 or 24 h). (Pea green and green lines vs light blue and blue lines, respectively in Figure 5A).

In addition, results confirmed that samples from high final pH group, at 0.5 h, had low levels of glycogen, since when plotting each spectrum only noise is observed (pea green line in Figure 5A); and a delta of absorbance was close to 10% of glycogen control (brown line in Figure 5A). Samples taken at 24 h from carcasses with normal pH presented a remarkable reduction in the relative intensities of absorbance, with respect to their basal control (0.5 hours) (light blue and blue lines, respectively, in Figure 5A).

## Discussion

It is known that AMPK is an enzyme activated in response to several stressors. Recently, a decrease in p-AMPK relative abundance in low pH with lactate conditions was shown in rat cardiomyocytes, as well as a trend towards increased phosphorylation with higher pH (Genders et al, 2019). Similarly, other authors showed that AMPK activity in cultured cardiomyocytes was suppressed upon acidic treatment but was stimulated upon alkaline treatment (Zhao et al 2016). Contrarily, but in *post-mortem* muscle, analyses of AMPK activity performed by FRET/spectrofluorometric assays, consistently showed a greater and sustained AMPK activity (400%) in muscle samples from carcasses categorized with normal final pH vs those with high pH (Apaoblaza et al, 2015). Here, a similar difference was found between both pH categories, 630 and 400% according to time evaluated at 0.5 and 24 h, respectively (Table 1).

Moreover, we showed that pH differences between both categories starting 120 minutes after entering the chamber and the differences between AMPK activity from both pH categories happened much earlier (before 30 minutes), in which the differences in pH there is not get (Galaz, 2014, Ramírez et al, 2013). On the other hand, AMPK activity did not appear to vary when pH declined by1 point (6.73 to 5.73, Table 1) or more in samples categorized with normal pH decline (7.76 to 5.71, Apaoblaza et al 2015).

Temperature and other stressors associated with cooling or freezing (ischemia and osmotic shock) affect the function of several tissues and organs in animals (Storey and Storey, 2004; Dalo et al, 1995; Hillman, 1998, Lutz and Reiners, 1997). Regarding AMPK activity regulation by environmental stressors both hypoxia and hypothermia in frogs increases AMPK phosphorylation level and AMPK activity (Bartrons et al., 2004). Furthermore, more than hypothermia freeze tolerance of *Rana sylvatica* revealed that during in vivo freezing, AMPK enzymes increased their activity (Storey, 2006). This suggests that AMPK activation would facilitate adaptation to hypothermia, freezing and thawing.

The decline in pH in skeletal muscles *post-mortem* is due to the formation of lactate from stored glycogen (Bendall, 1973). Furthermore, temperature linearly affects the rate of decline in muscle pH in *post-mortem* samples from body temperature down to 5 °C with a slower rate of decline at progressively lower temperatures (Cassens & Newbold, 1967). It is interesting that below 5 °C, the effect of temperature on the rate of pH decline is faster than at 5 °C (Cassens & Newbold, 1966). The primary enzymes that regulate the rate of *post-morterm* glycolysis in muscle are glycogen phosphorylase (GP) and phosphofructokinase (PFK) (Swatland, 1979). Effectively, previous works reported by Helmreich and Cori (1965) in living muscle from frog and by Newbold and Scopes (1967) in Ox muscle *post-mortem*, showed that of the 12 enzyme reactions involved in the conversion of glycogen into lactate; both the phosphorylase and phosphofructokinase reaction are of main importance in the regulation of glycogen breakdown in both conditions (living or *post-mortem*).

Newbold and Scopes (1967), suggested that the activation of glycolysis below 5°C appears to be due to the greater accumulation of AMP which stimulates glycogenolysis by activating GP b and stimulating PFK in *post-mortem* muscle. On the other hand, AMP also is the main and direct activator of AMPK and in *post-mortem* muscle, and has been showed to have a strong correlation between AMPK activation and pH decline, indicating that AMPK regulates glycolysis in *post-mortem* muscles (Du et al, 2007; Shen & Du, 2005; Shen et al., 2006). Glycolysis regulation by AMPK in *post-mortem* muscle is mediated, at least partially, through phosphorylation and activation of PFK-2, since fructose-2,6-diphosphate content, an allosteric activator of PFK-1, correlated well with AMPK activation and with glycolytic rate (Shen et al, 2007).

Few reports tackle *post-mortem* changes in AMPK activity associated to low temperature exposures. However, the management of carcasses in cooling regime, which varies according the plant, season, species, volume or intensity of the task. Studies in turkey muscles showed that high temperature early *post-mortem* could induce AMPK activation, which results in rapid glycolysis, thus affecting protein solubility and generating PSE characteristics (Zhu et al., 2013). High and sustained AMPK activity in *post-mortem* bovine muscle at low temperature, has been suggested as a new variable that, added to reserve level of glycogen, conditions the normal flow of glycogenolysis/glycolysis, required for normal transformation of muscle into meat. (Apaoblaza et al., 2015; Jerez-Timaure et al., 2019).

In the *post-mortem* management of carcasses, the “physiologic context” of the muscle include hypoxia and cooling. Regarding cooling, we showed that the carcases in cold chambers experience two cooling rates: The first (32 °C to 10 °C) near to −0.054°/min. The second, (10° to 2 °C), near to −0.015 °C/min. This study (Table 1) and other studies showed that AMPK activity, measured between 17-20 °C (room temperature) is 4 times higher in samples from carcasses with normal final pH than high final pH; and, the level of activity (in both categories) is the same at early (0,5 h) and after 24 h *post-mortem* (Apaoblaza et al, 2015) or is slightly higher (Jerez-Timaure et al 2019).

Here we showed that AMPK activity in *post-mortem* muscle is a parameter highly affected by the temperature of analysis, with maximal activity detected (both categories samples) at 17 °C (Figure 2 and 3). Effectively, AMPK activity in normal samples (0.5 h) evaluated at 17 ° C was nearly 90% higher than when evaluated at 25 or 37 ° C (Figure 2A and Figure 3). The same behaviour, although less pronounced, was observed from samples taken at 24 hours (near to 64%) (Figure 2B). We also showed that AMPK activity was totally suppressed when evaluation was carried out at 5 °C, a temperature that is reached approximately 12 hours from the entrance of carcasses to the cold chamber.

Interestingly, the optimal temperature (maximal AMPK activity) obtained in this study (17 °C), coincided with room temperature range (17-20 °C) used in the initial evaluation of the samples from this study (Table 1), and also the previous studies from our group (Apaoblaza et al 2015; Apaoblaza et al 2017; Jerez-Timaure 2019).

Newbold and Skopes (1967) pointed out that since the fall in pH is linearly related to lactate production, PFK and GP activities are reflected in final pH and can be assessed from changes in lactate or pH, and in hexose monophosphate concentration and total adenosine triphosphatase activity, respectively.

Those authors showed that PFK and GP activities before to 7 hour post-mortem, were not lower at 1 °C or 5 °C than at 15 °C; arguing that the decrease in metabolic rate expected as a result of lowering the temperature was counteracted by some factor(s) that accelerated the metabolism, as increase of GP step, evidenced by high hexose monophosphate concentration and high levels of AMP, a GP activator (Newbold and Skopes, 1967).

Interestingly, the pH time-course in temperature function from this study, the authors showed that in the muscle samples maintained at 15 °C the ultimate pH (24-48-72 h) is lower than those maintained to 1 or 5 °C (near to 0.4 pH point at 24 h); and these differences begin from 7-8 hours (Newbold and Scopes, 1967). Here, and also in previous studies (Ramirez et al 2013; Galaz, 2015), we observed that significant differences in the pH between both categories begin earlier than 7 hours, specifically 3 hours from the time that carcasses are put in the cold chamber, where the muscle temperature at this time is near to 20 °C (Figure 1).

Based on the final pH results, and analyses derived from Newbold and Scopes (1967), we argue that muscle samples from normal versus high pH, experience a more efficient and consistent glycogenolytic and glycolytic flow, from a higher initial glycogen reserve. This is because they already have a significant pH in delta at 3 hours, when the temperature is near to optimal temperature for AMPK activity

This may be the reason that in the study of Newbold and Scopes (1967), the sample muscles with similar initial conditions and maintained at 15 °C achieved a lower pH at 7, 24, 36 and 72 hours, than the samples muscles maintained at 1 °C and 5 ° C. Unfortunately, beyond the storage temperatures reported by Newbold and Scopes (1967), the time-course of temperature of these muscle samples was not recorded.

A relevant aspect associated with carcasses cooling, is that temperature decline of muscle in cold chambers vary depending on anatomical location (i.e. deep v. superficial muscles), the weight and fatness of the carcass, and temperature and air-speed conditions during chilling (Ferguson and Gerrard, 2014). Consequently, glycolytic rate varies not only between muscles, but also within a muscle (Tarrant and Mothersill 1977).

Regarding the latter, Tarrant and Mothersill (1977) showed that the rate of glycolysis in four beef muscles was, on average, 64% faster when the measurement was taken at a depth of 8 cm in the muscle compared with 5 cm. In this study, the authors point out the relationship between pH and *in situ* temperature variation. For example, low pH values (<6.0) coincided with high temperature (>30°C) in the muscles of round at 8 cm samples. Moreover, combinations of low temperature (<10°C) and high pH (>6.0) were are associated with cold-shortening (Tarrant and Mothersill,1977).

This finding suggests that the temperature of cooling applied in cold chamber, can be a critical control point of *post-mortem* glycogenolysis and glycolysis, and should allow the muscle piece (carcasses) to experience temperatures close to 15-17 °C, at less few hours, and not an accelerated cooling to 5 °C, capable of neutralizing AMPK activity and its efficient control of glycogenolytic/glycolytic flux. In addition to adenine nucleotides, glycogen and the non-physiological glycogen-mimic cyclodextrin may also regulate AMPK. However, while some authors have not detected an effect of glycogen on the activity of purified rat liver AMPK (Polekhina et al 2003), other authors report that glycogen and cyclodextrin inhibit the catalytic activity of recombinant AMPK and native rat liver AMPK (McBride et al., 2009). Similarly, it has been shown that glycogen regulation occurred both in the presence and in absence of AMP and was dependent on its binding to the carbohydrate-binding loop of the CBM (McBride et al, 2009; Scott et al, 2014). Interestingly, and related to AMPK/Glycogen interaction, McBride et al, (2009), showed that glycogen inhibits purified AMPK in cell-free assays, by direct action of GBD, and that it varied according to the branching content of the glycogen. Moreover, these authors showed that the oligosaccharides used, are allosteric inhibitors of AMPK that also inhibit phosphorylation and activation by upstream kinases, and suggesting that the GBD is a regulatory domain that allows AMPK to act as a glycogen sensor in vivo. Like the report of Polekhina et al. 2003, our results reveal that the enzymatic activity of AMPK was not affected by glycogen addition to sample homogenate from either of the two categories (Figure 4).

Effectively, with *in vitro* glycogen supplementation (60mmol / kg), we did not observe an inhibitory effect on AMPK activity, expected in samples with high initial and final activity. After supplementation reached more than double their initial concentration (136 mmol/kg), their activity remained unchanged. However, an activating effect was not observed, expected in the samples with low initial and final activity, which after supplementation reached a slightly higher concentration than the samples that showed high AMPK activity (76 mmol/kg).

Different results (final pH) were obtained in *post-mortem* pig muscle samples in which the high ultimate pH meat in oxidative muscle has been suggested to be caused by a lack of ante-mortem glycogen level. However, a recent study by England et al., 2016 showed that in the presence of excess glycogen and at 25°C, pig oxidative muscles produce a meat with a high final pH. These authors suggest that the porcine oxidative muscle (*in vivo* and *in vitro*) can and do stop *post-mortem* pH decline in presence of excess glycogen.

Glycogen levels estimated by relative absorbance (Figure 5A and 5B) confirm a lower availability of the polymer under high final pH conditions; since the glucose mmol reported at the initial time, (0.5 hours) are practically reduced to zero after 24 hours. The above situation is not observed in conditions of normal final pH, since although there is a reduction in amount of the polymer, it is possible to quantify it after 24 hours and recover it after ethanol precipitation with absorbance delta near to 0.5 AU, decreasing greater than 50% of the initial polymer after 24 hours.

For their part, results obtained after glycogen scavenging in the presence of iodide suggest that under conditions of high pH there is a lower basal amount of glycogen that drops close to 50% after 24 hours. Under conditions of acidic pH, basal glycogen content is substantially higher than in alkaline conditions, being almost 4 times higher, decaying at a reduction rate of close to 75% in 24 hours. In both conditions, there would be no abnormality in the type of polymer or the degree of ramifications thereof, behaving as a conventional glycogen and not as an amylopectin. This above suggests that phosphorylase and debranching activity operate correctly during the *rigor mortis* process and that observed muscle pH changes would not modulate these activities.

Our results give evidence that the *post-mortem* enzymatic activity of AMPK is conditioned by the metabolic context and signal transduction presents prior to animal slaughter. In the same context, Park et al. (2007), using regression analysis (lean meat colour and pH at 24 h) concluded that cattle with lower pH at 1 hour *post-mortem* has a faster decline in pH and resulted in lower pH at 24 h. In another work (Park et al., 2007), results could be understood as the high pH cattle (i.e., DFD condition) were resulted from their own initial potentiality prior to slaughter. Interestingly, the mean of pH registered at three hours in the study of were 6.6 and 6.0 0.6 pH point) for DFD and normal condition, respectively. In this work the mean initial pH at 0.5 hours were 6.91 and 6.73 (0.18 pH point) for high and normal pH categories, respectively. This difference is slightly greater than that recorded in the study of Apaoblaza et al. (2015) with 6.87 and 6.76 (0.11 pH point) at 0.5 hours and for the same categories.

Here, we report that the maximal AMPK activity in *post-mortem* mammalian muscle occurred close to 17 °C but not at 25 °C or 37 ° C, for other hand we showed that AMPK activity is close zero at 5 ° C. Interesting, AMPK activity has been suggested as an adaptation to hypothermia in animals evolutionarily remote. Effectively, hypothermia in frogs increases the AMPK phosphorylation levels and AMPK activity (Bartrons et al., 2004; Storey, 2006). Additionally, in sperm cells, we evaluated if pharmacological activation of AMPK improved the freezability of spermatozoa in stallions (Córdova et al., 2014) and showed that the AMPK in this cellular model did not respond to classical activators such as AMP, AICAR and metformin. We found that the AMPK activity in homogenate from stallion spermatozoa was 52% higher at 17 °C than at 5 °C (data non published) and we believe that the high cooling rates used in study of Cordova et al. (2014) between 20 °C to 5 °C (7.4 ° C / min and 1.8 °C / min in two slopes) was able to neutralize AMPK’s activity, since 5 ° C was reached in less than 10 minutes. Inherently, other authors have reported that porcine sperm stored at 17 °C (optimal temperature for commercial use of pig semen for several days) increased AMPK activity, longevity and sperm motility maintenance (Martin-Hidalgo et al, 2013).

Thermo-sensibility and thermo-stability of AMPK activity in muscle and other tissues and cellular types are is a field of study that can open interesting windows for improve the cooling managing of *post-mortem* muscle and also in other possible applications as a organs transplants and cell preservation.

## Conclusions

AMPK activity is highly sensitive to temperature, presenting maximum levels of activity at 17 °C and minimal levels at 5 ° C.

The *in vitro* excess of glycogen does not affect the enzymatic activity of AMPK in *post-mortem* cattle muscle.

Normal levels of pre-mortem muscle glycogen and AMPK activity determine the efficient glycolytic/glycolytic flow required for lactate accumulation and pH decline.

## Declarations

### Ethics approvals

Certificate No. 34-2011, Bioethics Committee for use of animals in research. Universidad Austral de Chile.

## Acknowledgements

The authors wish to thank Project FONDECYT 1120757, Chile, for financing this study, and FRIVAL slaughterhouse for providing the facilities for measurements.

## List of abbreviations

AMP: Adenosine monophosphate
AMPK: AMP-activated protein kinase
ATP: Adenosine triphosphate
GBD: Glycogen-binding domain
EF2: Elongation factor 2
AICAR: 5-Aminoimidazole-4-carboxamide ribonucleotide
mGS: Muscle glycogen synthase
CBM: Carbohydrate-binding module
PSE: Pale, soft and exudative
LD: Longissimus dorsi
T0: Temperature at 0-30 minutes
T24: Temperature at 24 hours
MGC: Muscle glycogen muscle
LA: Lactate
FRET: Fluorescence resonance energy transfer
GP: Glycogen phosphorylase
PFK: Phosphofructokinase
AU: Arbitrary units
DFD: Dark Firm Dry

## References

1. Hardie D, Hawley S and Scott J (2006) AMP-activated protein kinase-development of the energy sensor concept. J. Physiol 574(1). 7–15.

2. Hardie DG, Scott JW, Pan DA and Hudson ER (2003) Management of cellular energy by the AMP-activated protein kinase system. FEBS Lett 546, 113–120.

3. Kahn BB, Alquier T, Carling, D and Hardie DG (2005) AMP-activated protein kinase: ancient energy gauge provides clues to modern understand of metabolism. Cell Metab 1, 15–25.

4. Ramamurthy S and Ronnett GV (2006) Developing a head for energy sensing: AMP-activated protein kinase as a multifunctional metabolic sensor in the brain. J Physiol 574, 85–93.

5. Leff T (2003) AMP-activated protein kinase regulates gene expression by direct phosphorylation of nuclear proteins. Biochem Soc Trans 31, 224–227.

6. Hardie DG (2004) AMP-activated protein kinase: a master switch in glucose and lipid metabolism. Rev Endocr Metab Disord 5, 119–125.

7. Hallows KR (2005) Emerging role of AMP-activated protein kinase in coupling membrane transport to cellular metabolism. Curr Opin Nephrol Hypertens 1(4), 464–471.

8. Lizcano JM, Göransson O, Toth R, et al (2004) LKB1 is a master kinase that activates 13 protein kinases of the AMPK subfamily, including the MARK/PAR-1 kinases. EMBO J 23, 833–843.

9. Zhou G, Myers R, Moller DE et al (2001) Role of AMP-activated protein kinase in mechanism of metformin action. J Clin Invest 108, 1167–1174.

10. Hudson ER, Pan DA, James J, Lucocq JM, Hawley SA, Green KA, … & Hardie DG (2003) A novel domain in AMP-activated protein kinase causes glycogen storage bodies similar to those seen in hereditary cardiac arrhythmias. Current biology 13(10), 861–866.

11. Polekhina G, Gupta A, Michell BJ, et al. (2003) AMPK beta subunit targets metabolic stress sensing to glycogen. Curr Biol 13:867–871.

12. Storey JM, Storey KB. Cold hardiness and freeze tolerance. Functional metabolism: regulation and adaptation John Wiley and Sons, Inc. 2005 Feb 25:473–503.

13. Daló NL, Hackman JC, Storey KE, Davidoff RA (1995) Changes in motoneuron membrane potential and reflex activity induced by sudden cooling of isolated spinal cords: differences among cold-sensitive, cold-resistant and freeze-tolerant amphibian species. Journal of experimental biology 198(8), 1765–74.

14. Hillman SS (1988) Dehydrational effects on brain and cerebrospinal fluid electrolytes in two amphibians. Physiological zoology 61(3), 254–259.

15. Lutz PL, Reiners RA (1997) Survival of energy failure in the anoxic frog brain: delayed release of glutamate. Journal of experimental biology 200(22), 2913–2917.

16. Bartrons M, Ortega E, Obach M, Calvo MN, Navarro-Sabaté A, Bartrons R. (2004) Activation of AMP-dependent protein kinase by hypoxia and hypothermia in the liver of frog *Rana perezi*. Cryobiology 49(2): 190–194

17. Storey KB (2006) Reptile freeze tolerance: metabolism and gene expression, Cryobiology 52 1–16.

18. McBride A, Ghilagaber S, Nikolaev A, Hardie DG (2009) The glycogen-binding domain on the AMPK beta subunit allows the kinase to act as a glycogen sensor. Cell Metab; 9:23–34

19. Wojtaszewski JF, MacDonald C, Nielsen JN, Hellsten Y, Hardie DG, Kemp BE, … & Richter EA (2003) Regulation of 5′ AMP-activated protein kinase activity and substrate utilization in exercising human skeletal muscle. American Journal of Physiology-Endocrinology and Metabolism, 284(4), E813–E822.

20. Derave W, Ai H, Ihlemann J, Witters LA, Kristiansen S, Richter EA & Ploug T (2000) Dissociation of AMP-activated protein kinase activation and glucose transport in contracting slow-twitch muscle. Diabetes 49(8), 1281–1287.

21. Wojtaszewski JF, Jorgensen SB, Hellsten Y, Hardie DG and Richter EA (2002) Glycogen-dependent effects of 5-aminoimidazole-4-carboxamide (AICA)-riboside on AMP-activated protein kinase and glycogen synthase activities in rat skeletal muscle. Diabetes 51, 284–292.

22. Nielsen JN, Wojtaszewski JF, Haller RG, Hardie DG, Kemp BE, Richter EA & Vissing J (2002) Role of 5′ AMP-activated protein kinase in glycogen synthase activity and glucose utilization: insights from patients with McArdle’s disease. The Journal of physiology 541(3), 979–989.

23. Davies SP, Carling D & Hardie DG (1989) Tissue distribution of the AMP-activated protein kinase, and lack of activation by cyclic-AMP-dependent protein kinase, studied using a specific and sensitive peptide assay. European journal of biochemistry 186(1-2), 123–128.

24. Hardie DG & Sakamoto K (2006) AMPK: a key sensor of fuel and energy status in skeletal muscle. Physiology 21(1), 48–60.

25. Skurat AV, Wang Y, Roach PJ (1994) Rabbit skeletal muscle glycogen synthase expressed in COS cells. Identification of regulatory phosphorylation sites. Journal of Biological Chemistry 269(41), 25534–25542.

26. Jørgensen SB, Nielsen JN, Birk JB, Olsen GS, Viollet B, Andreelli F, … & Richter EA (2004) The α2–5′ AMP-activated protein kinase is a site 2 glycogen synthase kinase in skeletal muscle and is responsive to glucose loading. Diabetes, 53(12), 3074–3081.

27. Scott JW, Ling N, Issa SM, et al. (2014) Small molecule drug A-769662 and AMP synergistically activate naive AMPK independent of upstream kinase signaling. Chem Biol 21:619–627.

28. Du M, Shen QW, Zhu MJ (2005) Role of β-adrenoceptor signaling and AMP-activated protein kinase in glycolysis of postmortem skeletal muscle. Journal of agricultural and food chemistry 53(8), 3235–3239.

29. Shen Q and Du M (2005) Role of AMP-activated protein kinase in the glycolysis of postmortem muscle J Sci Food Agric 85:2401–2406.

30. Shen QW, Means WJ, Underwood KR, Thompson SA, Zhu MJ, McCormick RJ, Ford SP, Ellis M and Du M (2006) Early post-mortem AMP-activated protein kinase (AMPK) activation leads to phosphofructokinase-2 and-1 (PFK-2 and PFK-1) phosphorylation and the development of pale, soft, and exudative (PSE) conditions in porcine longissimus muscle. Journal of agricultural and food chemistry 54(15), pp.5583–5589.

31. Apaoblaza A, Galaz A, Strobel P, Ramírez-Reveco A, Jeréz-Timaure N & Gallo C (2015) Glycolytic potential and activity of adenosine monophosphate kinase (AMPK), glycogen phosphorylase (GP) and glycogen debranching enzyme (GDE) in steer carcasses with normal (< 5.8) or high (> 5.9) 24 h pH determined in M. longissimus dorsi. Meat Science 101, 83–89.

32. Chan T & Exton J (1976) A rapid method for the determination of glycogen content and radioactivity in small quantities of tissue or isolated hepatocytes. Analytical Biochemistry 71, 96–105.

33. Villarroel-Espíndola F, Tapia C, González-Stegmaier R, Concha II, & Slebe JC (2016) Polyglucosan molecules induce mitochondrial impairment and apoptosis in germ cells without affecting the integrity and functionality of sertoli cells. Journal of cellular physiology 231(10), 2142–2152.

34. Genders AJ, Martin SD, McGee SL & Bishop DJ (2019) A physiological drop in pH decreases mitochondrial respiration, and HDAC and Akt signaling, in L6 myocytes. American Journal of Physiology-Cell Physiology 316(3), C404–C414.

35. Zhao L, Cui L, Jiang X, Zhang J, Zhu M, Jia J, … & Huang Y (2016) Extracellular pH regulates autophagy via the AMPK–ULK1 pathway in rat cardiomyocytes. FEBS letters, 590(18), 3202–3212.

36. Galaz A (2014) Actividad de la enzima quinasa activada por AMP (AMPK) y enzimas glicogenolíticas en el pH final del músculo *Longissimus thoracis* en canales bovinas. Tesis de grado. Bioquímica. Universidad Austral de Chile.

37. Ramírez A, Galaz A, Strobel P, Apaoblaza A, Jerez N y Gallo C (2013) Actividad de la quinasa activada por adenosin monofosfato (AMPK) y su relación con las enzimas glicogenolíticas y el pH final del músculo *Longissimus dorsi* (LD) en canales de novillos del sur de Chile. XXIII Reunión de La Asociación Latinoamericana de Producción Animal (ALPA), 18 al 22 de noviembre, La Habana – Cuba.

38. Bendall JR. Postmortem changes in muscle. The structure and function of muscle. 1973;2(Part 1):243–309.

39. Cassens RG, Newbold RP (1967) Temperature dependence of pH changes in ox muscle post-mortem. Journal of Food Science 32, 13–14. doi:10.1111/j.1365-2621.1967.tb01947.x

40. Cassens RG & Newbold RP (1966) Effects of temperature on post-mortem metabolism in beef muscle. Journal of the Science of Food and Agriculture 17(6), 254–256.

41. Swatland HJ (1979) Low temperature activation ofpost mortem glycogenolysis in bovine skeletal muscle fibres. The Histochemical Journal 11(4), 391–398.

42. Helmreich E, Cori CF. Regulation of glycolysis in muscle. Advances in Enzyme Regulation. 1965 Jan 1;3:91–107.

43. Newbold RP, Scopes RK (1967) Post-mortem glycolysis in ox skeletal muscle. Effect of temperature on the concentrations of glycolytic intermediates and cofactors. Biochemical Journal 105(1), 127–136.

44. Zhu X, Ruusunen M, Gusella M, Ylä-Ajos M, Xu X, Zhou G & Puolanne E (2013) High early *post-mortem* temperature induces activation of AMP-activated protein kinase and development of pale, soft and exudative characteristics in turkey muscles. Meat Science, 93(3), 600–606.

45. Jerez-Timaure N, Gallo C, Ramírez-Reveco A, Greif G, Strobel P, Pedro AV & Morera FJ (2019). Early differential gene expression in beef Longissimus thoracis muscles from carcasses with normal (< 5.8) and high (> 5.9) ultimate pH. Meat Science, 153, 117–125.

46. Ferguson DM & Gerrard DE (2014) Regulation of *post-mortem* glycolysis in ruminant muscle. Animal Production Science 54(4), 464–481.

47. Tarrant PV, Mothersill C (1977) Glycolysis and associated changes in beef carcasses. Journal of the Science of Food and Agriculture 28, 739–749. doi:10.1002/jsfa.2740280813

48. England EM, Matarneh SK, Oliver EM, Apaoblaza A, Scheffler TL, Shi H & Gerrard DE (2016) Excess glycogen does not resolve high ultimate pH of oxidative muscle. Meat Science, 114, 95–102.

49. Park BY, Lee JM & Hwang IH (2007) Effect of postmortem metabolic rate on meat color. Asian-australasian journal of animal sciences 20(4), 598–604.

50. Córdova A, Strobel P, Vallejo A, Valenzuela P, Ulloa O, Burgos RA, … & Ramírez-Reveco A (2014). Use of hypometabolic TRIS extenders and high cooling rate refrigeration for cryopreservation of stallion sperm: presence and sensitivity of 5′ AMP-activated protein kinase (AMPK). Cryobiology, 69(3), 473–481.

51. Martin-Hidalgo D, de Llera AH, Yeste M, Gil MC, Bragado MJ & Garcia-Marin LJ (2013). Adenosine monophosphate-activated kinase, AMPK, is involved in the maintenance of the quality of extended boar semen during long-term storage. Theriogenology, 80(4), 285–294.

